# Genome-wide Expression Profiling and Pathway Analysis in Cyclic Stretched Human Trabecular Meshwork Cells

**DOI:** 10.1101/406082

**Authors:** Michelle D. Drewry, Jingwen Cai, Inas Helwa, Eric Hu, Sabrina Liu, Hongmei Mu, Yanzhong Hu, William M. Johnson, Pedro Gonzalez, W. Daniel Stamer, Yutao Liu

## Abstract

**Purpose:** Regulation of intraocular pressure is dependent upon homeostatic responses of trabecular meshwork (TM) cells to mechanical stretch. Despite the important roles of miRNAs in regulating TM function and aqueous outflow, it remains unclear how miRNA and their target genes interact in response to physiological cyclic mechanical stretch. We aimed to identify differentially expressed miRNAs and their potential targets in human TM cells in response to cyclic mechanical stress.

**Methods:** Monolayers of TM cells from non-glaucomatous donors (n=3-6) were cultured in the presence or absence of 15% mechanical stretch, 1 cycle/s, for 6 or 24-hours using computer-controlled Flexcell Unit. We profiled the expression of 800 miRNAs using NanoString Human miRNA assays and identified differentially expressed miRNAs using the Bioconductor Limma package. We identified differentially expressed genes using Operon Human Oligo Arrays with GeneSpring software. Pathway analysis with WebGestalt identified stretch-related pathways. We used Integrative miRNA Target Finder from Ingenuity Pathway Analysis to identify potential miRNA-mRNA regulations.

**Results:** We identified 540 unique genes and 74 miRNAs with differential expression in TM cells upon cyclic mechanical stretch. Pathway analysis indicated the significant enrichment of genes involved in Wnt-signaling, receptor protein serine/threonine kinase signaling, TGF-β pathway, and response to unfolded protein. We also identified several miRNA master regulators, including miR-19b-3p and miR-93-5p, which may act as switches to control several mechano-responsive genes.

**Conclusions:** This study suggests that cyclic mechanical stress of TM cells triggers alterations in the expression of both mRNAs and miRNAs implicated in glaucoma-associated pathways.

## Introduction

Glaucoma is a group of optic neuropathies with progressive loss of retinal ganglion cells, optic atrophy, and visual field loss (1, 2). Glaucoma affects more than 70 million people worldwide (3, 4). Primary open-angle glaucoma (POAG) is the most common type of glaucoma without an identifiable secondary cause and affects approximately 3 million Americans in the US with more than half of these patients unaware of their glaucoma condition (3, 5). Often, POAG remains undiagnosed or unrecognized by patients until visual field loss is clinically severe (1). Known risk factors for POAG include advanced aging, elevated intraocular pressure (IOP), a positive family history of glaucoma, and/or African/Hispanic ancestry (5, 6). Elevated IOP is the only modifiable risk factor in the clinic currently (7). Lowering IOP in glaucoma patients with pharmacological interventions or surgeries, can delay the progression of vision loss (8).

IOP is regulated by the secretion of the aqueous humor (AH) into the posterior chamber and its drainage from the anterior chamber of the eye (9, 10). AH is secreted from the ciliary epithelium into the posterior chamber, travels through the pupil, and exits the anterior chamber primarily through the conventional (trabecular meshwork or TM) pathway, with a small portion leaving via the unconventional pathway (9, 11). The balance between the rate of secretion and resistance to drainage determines the IOP. Excessive resistance to AH outflow through the TM pathway causes elevated IOP, which may lead to compression, deformation, and damage of the retinal ganglion cell axons (8). This elevated pressure will then produce a loss in peripheral vision, and if not treated, may result in total irreversible vision loss (8).

Even though dysfunction of the trabecular outflow pathway is responsible for elevated IOP, clinically, most of IOP-lowering medications target the unconventional outflow pathway or secretion. New medications targeting the predominant drainage pathway (the TM pathway) are in high demand, including two recently approved medications (8, 12, 13). Thus, rho kinase inhibitors and nitric oxide drugs are the results of research over the last 25 years that identified important regulatory processes in the trabecular outflow pathway (8).

Due to high levels of fluctuations in IOP from blinking, eye movement and ocular pulse, TM cells are constantly under mechanical forces and strain (9, 14, 15). TM cells react to this stress, by maintaining cellular and tissue homeostasis, and preventing injury (16, 17). Strain causes profound changes to cell morphology, affecting motility, stiffness, contraction, orientation, and cell alignment (14-16). Recent studies have indicated autophagy as one of the relevant stress response adaptive mechanisms (16, 18, 19). Other genes that are impacted by the mechanical strain include those involved in extracellular matrix synthesis/remodeling, cytoskeletal organization, and cell adhesion (16, 20-25). It remains unknown how these genes are regulated (16). Since miRNAs regulate gene expression, we aimed here to identify the miRNA profile changes in response to mechanical strain in TM cells and to determine how these specific miRNAs target the gene expression in the stretched TM cells.

Our study focuses on discovering specific pathways that may induce elevated IOP. We stretched cultures of human TM cells by 15% with 1 cycle/second and measured genome-wide gene and miRNA expression profiles and their respective signal pathways in response to the mechanical stretch. Our hypothesis for this study is that mechanical stress-responsive pathways in human TM are critical for maintaining AH outflow resistance homeostasis and thus IOP. The identification of such pathways will enable further study into finding therapeutic targets that regulate IOP in glaucoma patients.

## Methods

### Cell Culture

Within 48 hours post mortem, primary human TM cells were obtained from cadaver eyes without a history of eye disease (26, 27). Tissues were processed in accordance with the Declaration of Helsinki. Secondary cultures of cells expanded from primary isolates of human TM cells were grown at 37°C in 5% CO_2_ in low glucose Dulbecco’s Modified Eagle Medium (DMEM) with L-glutamine, 110 mg/ml sodium pyruvate, 10% fetal bovine serum – premium select (Atlanta Biologicals, Flowery Branch, GA, USA), 100 μM non-essential amino acids, 100 units/ml penicillin, and 100 μg/ml streptomycin from Invitrogen (ThermoFisher Scientific, Waltham, MA, USA), as previously described (26).

### Gene Expression and Analysis

Primary human TM cells from three non-glaucoma donors (n=3) were plated on collagen-coated flexible silicone bottom plates from Flexcell (Flexcell International Corporation, Burlington, NC, USA). After reaching confluence, TM cells were switched to serum-free DMEM for 3 hours, followed by cyclic mechanical stress for 6 hours (15% stretching, 1 cycle/second) using the computer-controlled FX-3000 Flexercell Strain Unit from Flexcell (25). Control cells were cultured under the same condition but without mechanical stress. Total RNA was extracted using the RNAeasy kit (Qiagen, Germantown, MD, USA) according to the recommended procedures by the manufacturer. Quality and quantity of the RNA were evaluated using Agilent Bioanalyzer 2100 (Agilent Technologies, Santa Clara, CA, USA). RNA from three stretched and three control TM cells were amplified and labeled with Cy5 dye, and a universal Reference human RNA from Stratagene, Agilent Technologies, was also labeled with Cy3 dye. Each TM sample and the reference RNA sample were hybridized to Operon Human Oligo Arrays version 3.0 (35k probes) at the Duke University Microarray Core Facility.

Raw data was normalized using LOWESS (locally weighted scatterplot smoothing), and the logRatio for each sample was downloaded from the GSE14768 dataset deposited in the NCBI GEO database (https://www.ncbi.nlm.nih.gov/geo/query/acc.cgi?acc=GSE14768) (25). We analyzed the expression data using the GeneSpring version 14.9 software from Agilent, performing the Welch’s t-test to derive the significance of differential expression and the fold change between stretched and non-stretch samples. Significantly differentially expressed genes were identified with a Benjamini-Hochberg (BH) false discovery rate (FDR) < 0.05 and a fold change ≥ 2 or ≤ −2. The list of genes was categorized using the online PANTHER function classification system (28). The list of the significantly differentially expressed genes was uploaded into the WEB-based Gene Set Analysis Toolkit (WebGestalt) for gene ontology and pathway analysis (29). The gene ontology analysis examined potentially enriched pathways or categories in biological processes, cellular components, and molecular function. These differentially expressed genes were also uploaded to Ingenuity Pathway Analysis (IPA) to search for potential upstream regulators.

### miRNA Expression and Analysis

Monolayers of human TM cells from six donors (n=6) were maintained in low glucose DMEM media with 10% FBS and underwent cyclic mechanical stretch (15%, 1 cycle/second) for 24 hours, as described previously (30). Controls were under the same condition for 24 hours but without the cyclic mechanical stretch. Total RNA was extracted using the mirVana miRNA isolation kit from Thermo Fisher Scientific. RNA quantity was measured using the Tecan microplate reader M200 (Tecan Group Ltd., Mannedorf, Zurich, Switzerland), and RNA quality was evaluated using the Agilent Bioanalyzer 2100 with RNA Pico Chips. 100 ng of total RNA was used to measure the expression of 800 pre-selected human miRNAs using the nCounter Human v3 miRNA expression assay kit from NanoString Technologies (Seattle, WA, USA), as previously described (31). After specific sample preparation and overnight hybridization, digital readouts of the relative miRNA abundance were obtained, translating to miRNA expression.

The raw data RCC files produced by nCounter were processed with the NanoString nSolver 3.0 software using a previously described analysis pipeline (31). Each sample was run through several quality control checks before analysis. These checks examined the quality of imaging, binding density, positive control linearity, and positive control limit of detection. Based on RNA content, the raw miRNA counts were normalized using the trimmed geometric mean with the nSolver software. The geometric mean calculated the average number of counts for each sample using the miRNAs with the median 40% of the counts. This particular technique is similar to the Trimmed Mean of M method, which was shown to be successful in normalizing miRNA counts (32). To account for technology-associated sources of variation, the miRNA counts were normalized further using the geometric mean of the positive control probes.

After normalization, the data was exported from nSolver into CSV files and imported into the R Language and Environment for Statistical Computing. Using NanoString’s recommended methods, the background level probes for the control and stretched samples were identified within R. The negative control probes for each sample were extracted and multiplied by the corresponding normalization factors produced by nSolver. A Welch’s t-test was performed for each probe to compare the normalized negative control counts to the sample counts. If the counts for a probe were not significantly different than the negative controls (p>0.05), then it was considered background. Finally, the limma Bioconductor package was used to perform the differential analysis with paired samples after the background level probes were identified for each sample type. The differentially expressed probes were those with an absolute log2 fold change ≥ 1.5, a p-value ≤ 0.05, and an absolute difference in counts ≥ 5. Since the laboratory culture condition may change the miRNA expression in the cultures of human TM cells isolated from TM tissues, we examined whether these differentially expressed miRNAs were present in human TM tissues using our published miRNA expression profile from seven non-glaucoma human TM tissues with miRNA-Seq (33).

### Integrative Analysis of miRNA and mRNAs

miRNAs are known to regulate mRNA expression of many target genes. One miRNA may target many genes, and one gene could be regulated by many miRNAs. To identify miRNAs that potentially regulate the genes differentially expressed during the cyclic mechanical stretch of human TM cells, we used the miRNA Target Filter function in the Ingenuity Pathway Analysis (IPA, Qiagen). Briefly, we uploaded the list of differentially expressed miRNAs with expression fold changes and the list of differentially expressed mRNAs with expression fold changes to the IPA. The miRNA Target Filter in IPA detects target genes from multiple sources (TargetScan, TarBase, miRecords, and Ingenuity Knowledge Base) (34-36) and identifies which genes in the uploaded mRNA gene list are potential targets of the selected miRNAs. We required that the expression of interested miRNAs be negatively paired with their potential mRNA targets. If the miRNA expression is increased, then the expression of its target genes is decreased and vice versa. The target genes could include those experimentally validated in the literature or those predicted with high or moderate confidence. For accuracy, we limited our miRNA targets to those experimentally validated.

Although the cultures of human TM cells were derived from human trabecular meshwork tissues, gene expression profile may change due to laboratory culture conditions. Using the miRNA expression data from eight non-glaucomatous human TM tissue, we examined whether these specific miRNAs in TM cells were expressed in human TM tissue (33). Their expression in normal human TM tissues might suggest their potential physiological role in aqueous outflow dynamics.

### Availability of Data and Materials

The miRNA dataset supporting the conclusions of this article is available in the NCBI Gene Expression Omnibus and are accessible through the GEO Series accession number GSE113755 (https://www.ncbi.nlm.nih.gov/geo/query/acc.cgi?acc=GSE113755).

## Results

### mRNA Differential Expression Analysis

From gene differential analysis, using an absolute fold change ≥2 and a FDR value ≤0.05, we identified a total of 540 unique genes with significant differential expression (**Supplemental Table S1**). Of these, 276 genes were down-regulated and 264 genes up-regulated in the mechanically stretched human TM cells compared to non-stretched human TM cells. We have selected the top 20 down-regulated and top 20 up-regulated genes with the largest fold changes, listed in Table 1. Using the online PANTHER classification system, we found that the top functional categories of these genes were related to binding, catalytic activity, structural molecule activity, and transporter activity. The main classes of these proteins were related to nucleic acid binding, transcription factor, enzyme modulator, hydrolase, transferase, and cytoskeletal proteins. The top related pathways included gonadotropin-releasing hormone receptor pathway, CCKR signaling, apoptosis pathway, angiogenesis, Wnt-signaling pathway, integrin-signaling, and inflammation.

**Table 1.** List of differentially expressed genes in primary human TM cells in response to cyclic mechanical stretch (15%, 1 cycle/s, 6 hrs).

| Gene Symbol | Accession # | Fold Change | FDR P-value |
| --- | --- | --- | --- |
| *DHRS9* | NM_005771 | 1086.39 | 0.02 |
| *ESM1* | NM_007036 | 33.41 | 0.02 |
| *SCML4* | AK074125 | 29.74 | 0.02 |
| *HOXC13* | NM_017410 | 25.02 | 0.04 |
| *BMP2* | NM_001200 | 20.06 | 0.02 |
| *NP25* | NM_013259 | 19.73 | 0.02 |
| *PTHLH* | NM_002820 | 16.73 | 0.03 |
| *EVA1A* | NM_032181 | 12.27 | 0.01 |
| *PTX3* | NM_002852 | 9.77 | 0.02 |
| *MMP1* | NM_002421 | 8.85 | 0.02 |
| *GPAT3* | NM_032717 | 8.32 | 0.02 |
| *HSPA1B* | NM_005346 | 8.12 | 0.02 |
| *KCNC4* | NM_004978 | 7.97 | 0.02 |
| *MFSD2A* | AK093223 | 7.67 | 0.04 |
| *PNP* | NM_000270 | 7.61 | 0.01 |
| *PPP1R15A* | NM_014330 | 7.11 | 0.02 |
| *RGS17* | NM_012419 | 6.96 | 0.04 |
| *CHST11* | NM_018413 | 6.80 | 0.03 |
| *ARL7* | NM_005737 | 6.47 | 0.02 |
| *DKK1* | NM_012242 | 6.46 | 0.01 |
| *BTEB1* | NM_001206 | -13.88 | 0.03 |
| *CNTN3* | AB040929 | -13.41 | 0.04 |
| *NR4A2* | NM_006186 | -13.28 | 0.02 |
| *NR4A1* | NM_173158 | -12.59 | 0.03 |
| *NR4A1* | NM_173158 | -10.82 | 0.02 |
| *FAM83D* | NM_030919 | -10.39 | 0.02 |
| *DLX2* | NM_004405 | -9.82 | 0.01 |
| *EGR3* | NM_004430 | -8.32 | 0.02 |
| *C6orf111* | NM_032870 | -7.02 | 0.03 |
| *TNFSF10* | NM_003810 | -6.62 | 0.02 |
| *FBXO32* | NM_058229 | -6.56 | 0.02 |
| *SLC40A1* | NM_014585 | -6.41 | 0.02 |
| *PDE5A* | NM_001083 | -6.26 | 0.03 |
| *ZFP36L2* | NM_006887 | -5.68 | 0.02 |
| *CEMIP* | AB033025 | -5.37 | 0.05 |
| *TIEG* | NM_005655 | -5.25 | 0.02 |
| *RASL11B* | NM_023940 | -4.97 | 0.03 |
| *ATF3* | NM_004024 | -4.89 | 0.02 |
| *ZNF331* | NM_018555 | -4.83 | 0.03 |
| *GUCY1A3* | NM_000856 | -4.74 | 0.04 |

To identify enriched functional categories in the differentially expressed mRNAs via gene ontology, the list of these 540 genes was uploaded to WebGestalt. A number of these genes were identified as being significantly involved in many biological processes, such as regulation of transmembrane receptor protein serine/threonine kinase signaling pathway, canonical Wnt receptor signaling pathway, response to unfolded protein, and ribosome biogenesis (**Supplemental Figure S1**). Pathway analysis based on KEGG Pathway database (Kyoto Encyclopedia of Genes and Genomes) indicated the following enriched KEGG pathways in response to cyclic mechanic stretch: ribosome biogenesis, MAPK signaling, protein processing in ER (endoplasmic reticulum), metabolic pathways, P53 signaling pathway, and TGF-β signaling pathway (**Supplemental Table S2**). Ingenuity Pathway Analysis indicated that the top upstream regulators to the differentially expressed genes included *TGF-β1, HDAC, PDGF-BB*, and *MYC*.

### miRNA Differential Analysis

Using NanoString nCounter Human miRNA Assay for 800 pre-selected human miRNAs, we detected the expression of 373 different miRNAs in the cultures of human TM cells (**Supplemental Table S3**). Using a p-value cutoff < 0.05 and absolute fold change > 1.5, 74 miRNAs (10 down- and 64 up-regulated) were differentially expressed in cyclic mechanically stretched TM cells (Table 2). The top most significantly expressed miRNAs were miR-32-5p, miR-4286, miR-135-5p, miR-136-5p, miR-93-5p, miR-19b-3p, and miR-27a-3p (p value = 1.2×10^−5^, 1.3×10^−5^, 4.5×10^−5^, 7.0×10^−5^, 2.8×10^−4^, 3.1×10^−4^, and 9.1×10^−4^, respectively). The miRNAs with the greatest fold change in response to stretch included miR-4286, miR-32-5p, miR-136-5p, miR-93-5p, miR-135-5p, miR-21-5p, and miR-19b-3p, with fold changes ranging from 1.98 to 2.42. Consistent with previous reports, miR-100-5p, miR-27a-3p, miR-27b-3p, miR-24-3p, miR-16-5p, let-7f-5p, and let-7i-5p were significantly upregulated in stretched versus control TM cells (37). IPA showed that these miRNAs are mostly associated with cellular functions related to proliferation, development, cell cycle regulation, and cellular movement. We found that 29 of these 74 miRNAs were expressed in non-glaucomatous human trabecular meshwork tissues with normalized sequencing counts ≥ 10 (Figure 1).

**Table 2.** List of differentially expressed miRNAs in primary human TM cells in response to the cyclic mechanical stretch (15%, 1 cycle/s, 24 hrs)

| miRNA ID | Fold Change | P-value | miRNA ID | Fold Change | P-value |
| --- | --- | --- | --- | --- | --- |
| miR-32-5p | 2.38 | 1.2E-05 | miR-137 | 1.59 | 1.3E-02 |
| miR-4286 | 2.42 | 1.3E-05 | miR-31-5p | 1.59 | 1.4E-02 |
| miR-135b-5p | 2.03 | 4.6E-05 | miR-455-5p | 1.57 | 1.4E-02 |
| miR-136-5p | 2.11 | 7.0E-05 | miR-152-3p | 1.54 | 1.6E-02 |
| miR-19b-3p | 1.98 | 3.1E-04 | miR-24-3p | 1.67 | 1.6E-02 |
| miR-93-5p | 2.07 | 2.8E-04 | miR-30a-5p | 1.68 | 1.5E-02 |
| miR-27a-3p | 1.68 | 9.1E-04 | miR-3151-5p | -1.50 | 1.6E-02 |
| miR-362-3p | 1.68 | 1.3E-03 | miR-411-5p | 1.50 | 1.6E-02 |
| miR-151a-3p | 1.78 | 1.6E-03 | miR-26b-5p | 1.57 | 1.7E-02 |
| miR-25-3p | 1.94 | 1.7E-03 | miR-100-5p | 1.82 | 1.7E-02 |
| miR-337-5p | 1.72 | 1.8E-03 | miR-221-5p | 1.50 | 1.8E-02 |
| miR-140-5p | 1.82 | 2.6E-03 | miR-628-3p | 1.49 | 1.9E-02 |
| miR-497-5p | 1.57 | 2.9E-03 | miR-1290 | -1.43 | 2.0E-02 |
| miR-20a/b-5p | 1.70 | 3.3E-03 | miR-376c-3p | 1.49 | 2.1E-02 |
| miR-106a-5p/17-5p | 1.66 | 4.3E-03 | miR-642a-5p | 1.45 | 2.1E-02 |
| miR-15a-5p | 1.85 | 4.7E-03 | miR-185-5p | 1.45 | 2.2E-02 |
| miR-27b-3p | 1.92 | 4.0E-03 | miR-28-5p | 1.54 | 2.6E-02 |
| miR-374a-5p | 1.93 | 4.5E-03 | let-7i-5p | 1.72 | 2.8E-02 |
| miR-188-5p | -1.52 | 5.1E-03 | miR-16-5p | 1.64 | 2.8E-02 |
| miR-29a-3p | 1.96 | 5.9E-03 | miR-29c-3p | 1.69 | 2.8E-02 |
| miR-21-5p | 1.98 | 6.4E-03 | miR-301a-3p | 1.47 | 2.9E-02 |
| miR-127-3p | 1.67 | 9.9E-03 | miR-542-3p | 1.43 | 3.0E-02 |
| miR-1275 | -1.54 | 1.1E-02 | miR-1206 | 1.44 | 3.3E-02 |
| miR-187-3p | -1.52 | 1.0E-02 | miR-574-3p | 1.59 | 3.3E-02 |
| miR-190a-5p | 1.52 | 8.7E-03 | miR-590-5p | 1.40 | 3.4E-02 |
| miR-1972 | -1.50 | 8.0E-03 | miR-409-3p | 1.42 | 3.6E-02 |
| miR-199a-5p | 1.73 | 9.0E-03 | miR-505-3p | 1.41 | 3.7E-02 |
| miR-2117 | 1.49 | 9.4E-03 | miR-34a-5p | 1.60 | 4.0E-02 |
| miR-218-5p | 1.63 | 9.1E-03 | miR-1827 | 1.42 | 4.0E-02 |
| miR-33a-5p | 1.48 | 1.1E-02 | miR-520f-3p | -1.40 | 4.2E-02 |
| miR-376a-3p | 1.63 | 1.1E-02 | let-7f-5p | 1.49 | 4.2E-02 |
| miR-494-3p | 1.77 | 1.0E-02 | miR-639 | 1.42 | 4.3E-02 |
| miR-495-3p | 1.53 | 1.1E-02 | miR-125a-5p | 1.62 | 4.5E-02 |
| miR-642a-3p | 1.49 | 7.4E-03 | miR-566 | -1.30 | 4.7E-02 |
| miR-98-5p | 1.70 | 9.0E-03 | miR-147b | -1.39 | 4.7E-02 |
| miR-1323 | -1.47 | 1.2E-02 | miR-323a-3p | 1.38 | 4.9E-02 |
| miR-19a-3p | 1.55 | 1.2E-02 | miR-340-5p | 1.40 | 4.9E-02 |

**Figure 1.**
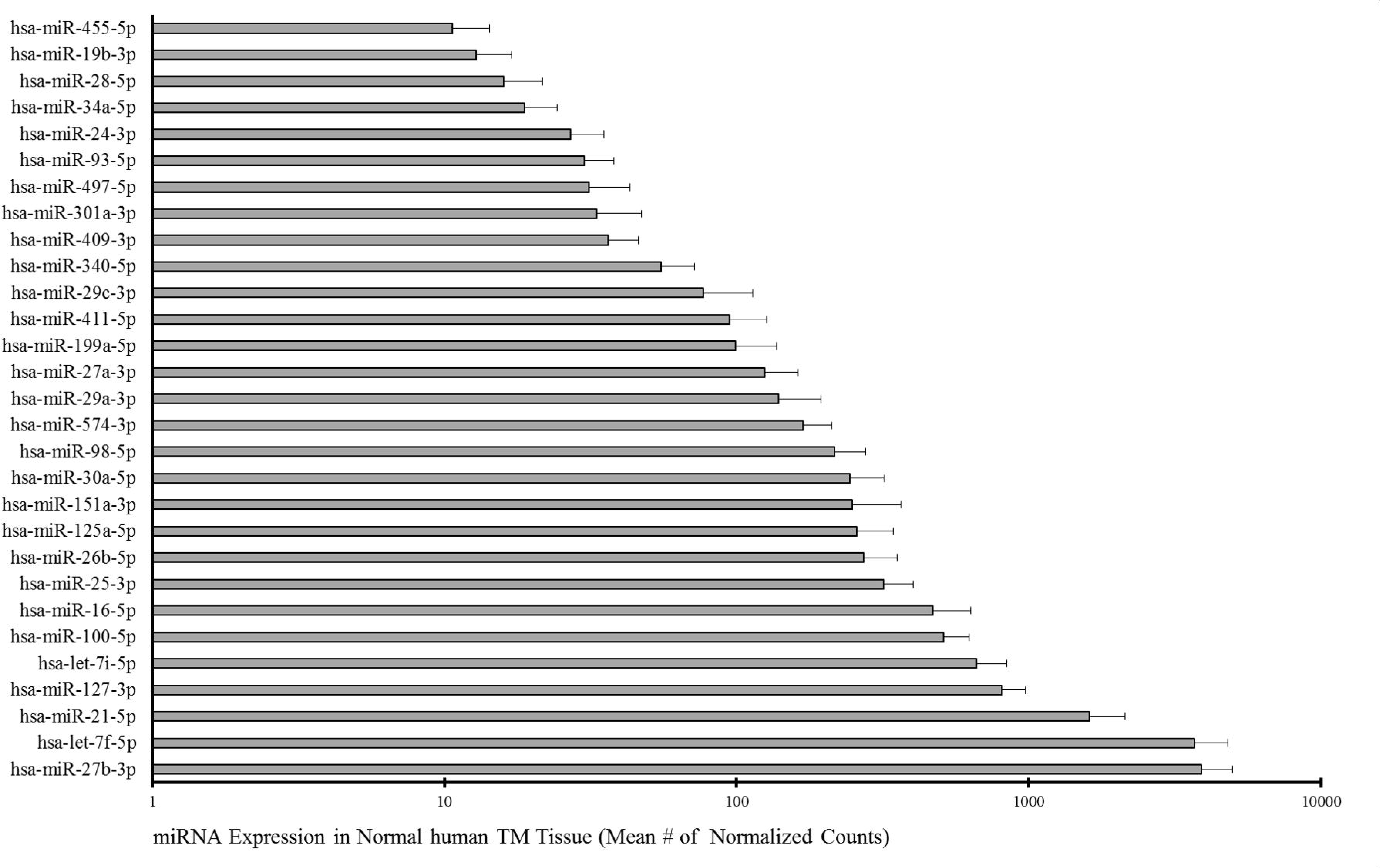
Expression level of cyclic mechanic stretch-responsive miRNAs in non-glaucoma human TM tissue (n=6). Error bars represent SEM.

### Integrative Analysis of miRNA and mRNAs

Using the miRNA Target Finder in IPA and restricting the analysis to only experimentally validated miRNA targets, we identified 8 unique miRNAs that target 17 mRNA genes in response to cyclic mechanical stretch in the cultures of human TM cells (Table 3). Most of these miRNAs targeted more than one gene, and several genes were targeted with more than one miRNAs (Figure 2). For example, increased expression of miR-15-5p inhibited the expression of *NT5DC1, OGT, OSGEPL1, RHOT1, SLC35A1*, and *WEE1*, while increased expression of miR-93-5p inhibited the expression of *BAMBI, HBP1, MAP3K12*, and *MYLIP*. The reduced expression of *MYLIP* was associated with the increased expression of miR-32-5p, miR-93-5p, and miR-19b-3p. The reduced expression of *WEE1* was related to the increased expression of miR-15a-5p and miR-27b-3p. Using our previous data on miRNA expression profile with 7 non-glaucomatous TM tissues (33), we found that six miRNAs (miR-125a-5p, miR-27b-3p, miR-93-5p, miR-29a-3p, miR-19b-3p, and miR-30a-5p), but not miR-15a-5p or miR-32-5p, were expressed in human TM tissue. This integrative miRNA target analysis identified potential regulatory expression networks in response to the cyclic mechanical stress in human primary TM cells.

**Table 3.**
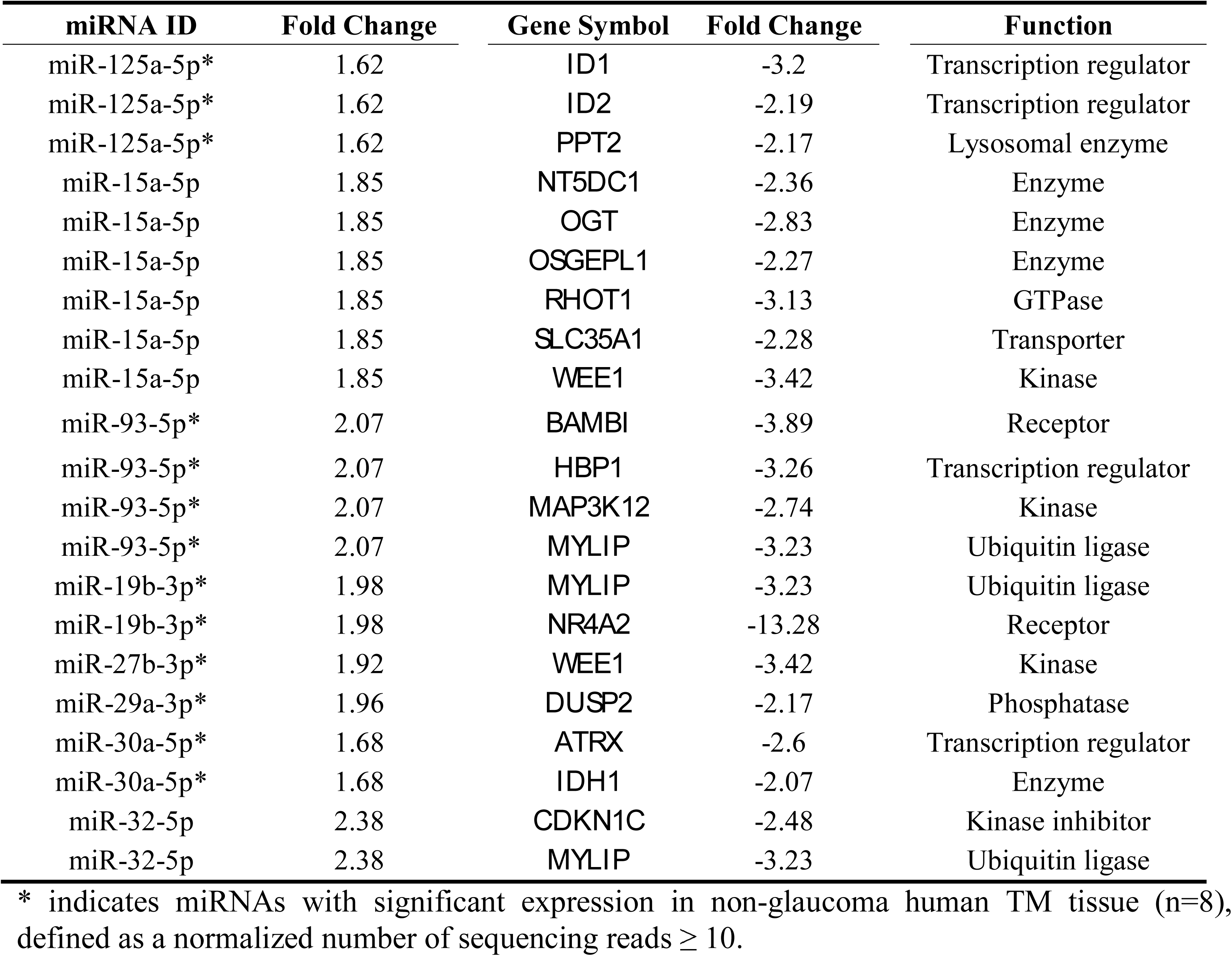
List of the interacting miRNAs and mRNA target genes in response to the cyclic mechanical stress in primary human TM cells

**Figure 2.**
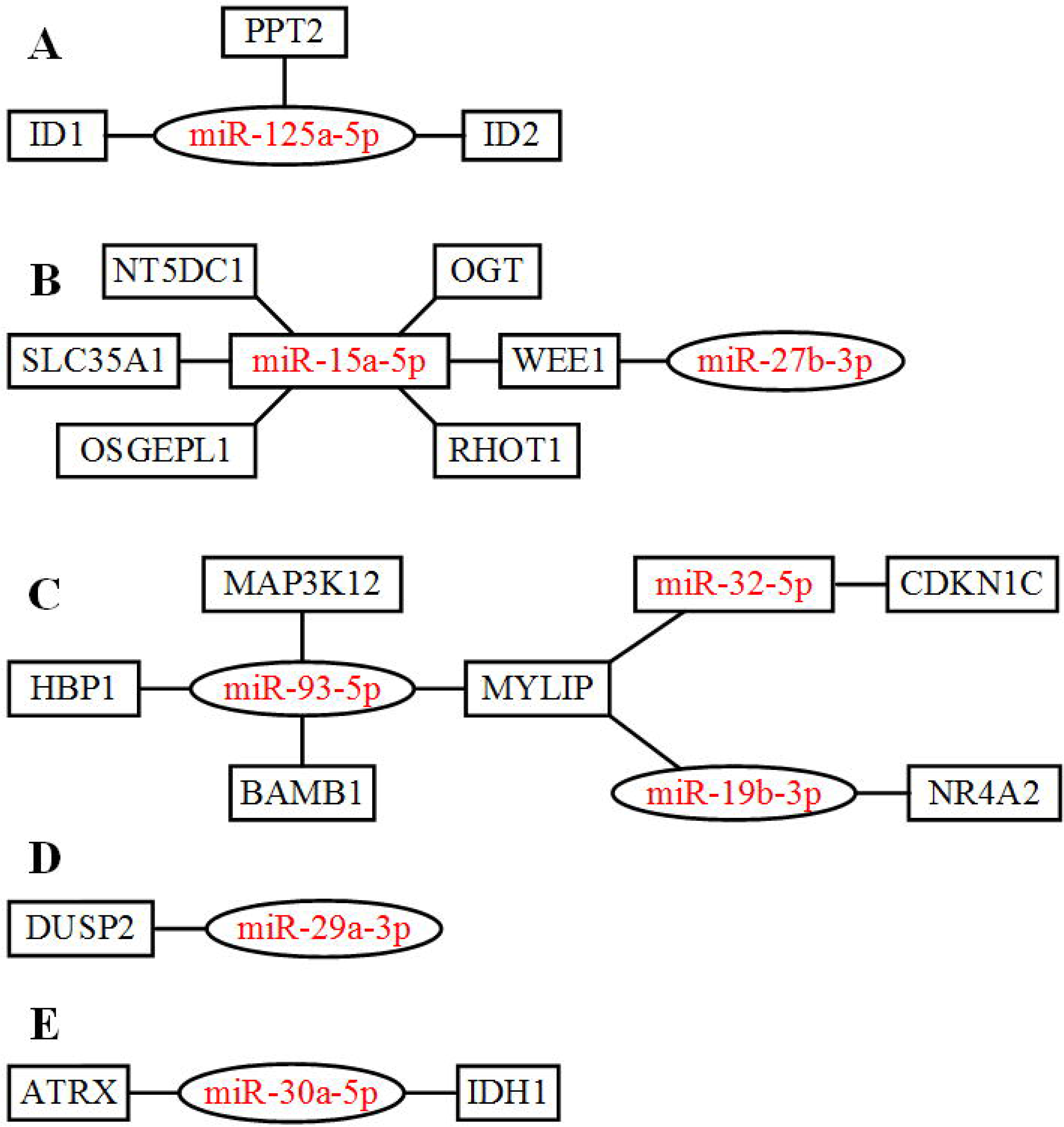
Gene network of cyclic mechanic stretch-responsive miRNAs, denoted in red, and their validated target genes. The miRNAs within an oval have significant expression in non-glaucoma human TM tissues (n=8), with ≥ 10 normalized sequencing reads.

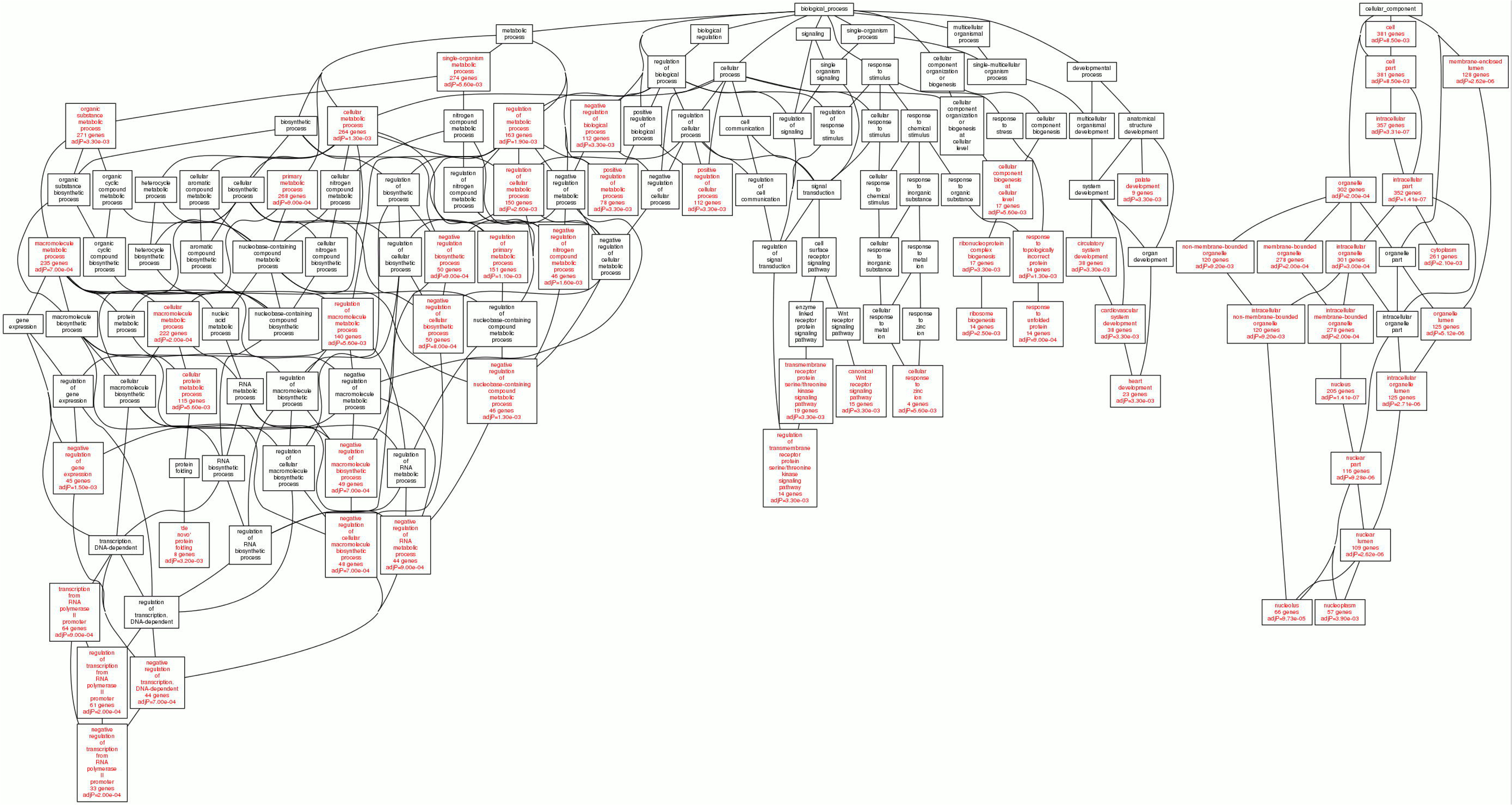

## Discussion

### Overview

We have identified 540 genes and 74 miRNAs differentially expressed in primary human trabecular meshwork cells in response to cyclic mechanical stress. Pathway analysis using these genes and miRNAs indicated potential involvement of Wnt receptor signaling pathway, unfolded protein response, serine/threonine kinase signaling pathway, TGF-β signaling pathway, mTOR signaling pathway, and apoptosis pathway. Our miRNA analysis suggests alterations in miRNAs potentially related to cell proliferation, development, cell cycle regulation, and cellular movement. Although the main findings of the expression responses to cyclic mechanical stress are similar to other mechanotranduction studies (22, 38), this is the first time that the Wnt-signaling pathway, unfolded protein response, and the serine/threonine kinase signaling pathway have been associated with cyclic mechanical stretch in monolayers of human TM cells.

### Wnt-signaling

Our analysis indicated significant enrichment of the canonical Wnt receptor signaling pathway (15 differentially expressed genes) in response to cyclic mechanical stretch in primary TM cells. Many studies have identified the activation of Wnt signaling as a normal physiological response to mechanical loading in bone and periodontal tissue (39). For example, Wnt signaling has been demonstrated to act downstream of mechanical activity and is required for joint patterning and chondrocyte maturation (40). Interestingly, the response of Wnt signaling to cyclic mechanical stress could be impaired by the aging process, i.e. the response of Wnt signaling to the mechanical stress is age-dependent, with a previous study showing more response in young adult mice (5-12 month old) and less or no response in old adult mice (22-month old) (41). Wnt inhibitor can induce the increased cellular stiffness in human trabecular meshwork cells (9), which might be related to the glaucoma progression (42). Wnt is present in the outflow facility as a local mediator (9). Importantly, an antagonist to Wnt signaling, sFRP-1increases outflow facility in perfused human anterior segments, and the overexpression of sFRP-1 leads to elevated IOP in mice (11, 43). Together this evidence suggests a homeostatic role for Wnt signaling in the trabecular outflow pathway and in regulation of IOP. Future studies will be needed to examine how this mechanical stress-induced Wnt signaling functions in glaucomatous TM cells or aged human TM cells. The Wnt signaling pathway could be regulated by several differentially expressed miRNAs, including miR-374a (44), miR-34a targeting p53 (45), let-7, and miR-31-5p (46). In summary, the interaction of Wnt signaling and miRNA regulation clearly contributes to the molecular response of human TM cells to cyclic mechanical stress.

### Unfolded protein response/ER stress

Unfolded protein response and related ER (endoplasmic reticulum) stress can be induced by cyclic mechanical stretching in smooth muscle cells and cardiomyocytes (47, 48), in which ER stress can lead to apoptosis and inflammation (49). ER stress and unfolded protein response have been implicated in the pathogenesis of glaucoma through the failure to eliminate misfolded/damaged proteins, leading to functional impairment of TM cells and increased outflow resistance (49, 50). Reducing ER stress and unfolded protein response in TM cells chemically can prevent dexamethasone-induced ocular hypertension in mice (49). Rare variants in key regulators of unfolded protein response (*ACADVL, EXOSC3, KDELR3, SHC1, SRPRB, SYVN1, TATDN2*, and *TPP1*) were found to be significantly enriched in POAG patients with elevated IOP (51). Differentially expressed miRNAs miR-455-5p, miR-34a-5p, miR-199a-5p, miR-221-5p, miR-494-3p, miR-29a-3p, miR-93-5p, miR-25-3p, and miR-30a have been known to target the regulators of unfolded protein response (52, 53). This evidence in conjunction with the differential expression of mRNAs and miRNAs associated with unfolded protein response provide additional support for its potential contribution to IOP regulation through TM cells. Further investigation into how unfolded protein response is damaged in glaucomatous TM cells from human donors will be necessary.

### Receptor serine/threonine kinase pathway/TGF-β Pathway

Receptor serine/threonine kinase pathway are mediated by the TGF-β type I and type II receptors (54). Cyclic mechanical stress has been shown to potentially activate TGF-β/BMP pathway through the SMAD-mediated Runx2 pathway or the non-SMAD-mediated MAPK pathway (55, 56). TGF-β and its related pathways have been strongly implicated in the pathogenesis of glaucoma and IOP regulation (54, 57-60). TGF-β may increase outflow resistance by altering extracellular matrix homeostasis and cell contractility in the TM through interactions with other proteins and signaling molecules (54, 58, 60). TGF-β treatment results in the regulation of many miRNAs, including the up-regulation of miR-21, miR-181, miR-494, miR-10b, miR-27a, miR-183, miR-182, miR-155, and miR-451 and the down-regulation of miR-200, miR-34a, miR-203, miR-584, and miR-450b-5p (61). Most members of TGF-β pathway may be targeted by a number of miRNAs, including miR-18a, miR-24, let-7, miR-744, miR-30, miR-200, miR-128a, miR-21, miR-17, miR-148a, miR-99a/b, and miR-92b (61, 62). Our study identified the differential expression of miR-24-3p, miR-27a, miR-21, miR-4286, and miR-29a, which all potentially target known genes involved in TGF-β pathway-mediated extracellular matrix homeostasis in TM cells.

### Conclusions and Future Work

Our study not only identified new miRNAs related to cyclic mechanical stretch in TM cells, but also replicated many previously reported miRNAs involved in this process. Several of our identified miRNAs, including miR-16, miR-27a, miR-27b, let-7f, miR-26a, miR-24 and miR-7, have been reported to be differentially expressed in TM cells after 3-hour cyclic mechanical stretching (63). miR-151 can regulate the expression of MMP13 (64), and miR-21 may regulate the expression of MMP9 and TIMP-1 (65). Both MMPs and TIMPs have been implicated in outflow resistance (66). An IOP-and POAG-associated gene *ABCA1* may be regulated by two differentially expressed miRNAs: miR-135 and miR-33a (67-69). Current IOP lowering therapies focus on reducing IOP without addressing extracellular matrix homeostasis processes in the TM (11). More studies would be beneficial to understand the role of TGF-β in POAG at the levels of production, activation, downstream signaling, and homeostatic regulation (58).

Our integrative miRNA-mRNA expression analysis identified several miRNA-mRNA regulatory expression interactions in the TM cells, highlighting several miRNA master regulators, such as miR-125a-5p, miR-15a-5p, miR-93-5p, miR-19b, miR-30a, and miR-32-5p. miR-15a has been reported to be down-regulated in stress-induced senescent human TM cells (70). miR-93 was found to have increased expression in glaucomatous TM cells and induced apoptosis in these TM cells (71). Since our integrative analysis was restricted to only those experimentally validated miRNA targets, additional master miRNA regulators may be identified if we expanded our analysis to include predicted targets with high or moderate confidence, though this would potentially increase the number of false positive findings.

Despite the strength of our experimental and bioinformatics analyses, our study has several limitations. First, the sample size of primary human TM cells was limited, derived from cells isolated from only 3 or 6 donor eyes. It would be better to have more donor-derived TM cells from both males and females, with similar ethnical background and similar age distributions. Second, the cyclic mechanical stretch experiment was done at single time point with 6 or 24 hours. Including a time-series design with many different time points would be beneficial in identifying time-dependent molecular responses, and it would be better to have the expression of miRNAs and mRNAs done at the same time point. Third, all the TM cells used in our study were derived from unaffected non-glaucomatous postmortem donors. Including TM cells derived from glaucoma-affected postmortem donors will be necessary to further explore mechanical stress-induced changes in miRNA and mRNA expression. These factors will be considered and included in our future experiments.

In summary, our genome-wide mRNA and miRNA expression profiling has identified a large number of differentially expressed genes and miRNAs in response to cyclic mechanical stretch in primary human TM cells. Our analysis has identified several important signaling pathways involved in this response, such as TGF-β, Wnt signaling, and ER stress. The miRNA-mRNA integrative analysis found several miRNA master regulators, suggesting their potential role in TM cellular function in response to cyclic mechanical stress. More functional studies are needed to further validate the role of these miRNAs and mRNAs in relation to TM cellular function.

## Acknowledgements

We thank the financial support from The Glaucoma Foundation, The Glaucoma Research Foundation, The BrightFocus Foundation, NIH R01EY023242, R21EY028671, R01EY022359, and R01EY023287. Financial support from Fight for Sight is gratefully acknowledged. The funding sources have no influences on the experimental design, data generation and analysis. We sincerely thank all the donors for their ocular samples. This study would not be feasible without these precious samples. This manuscript has been presented at the ISER/BrightFocus 2017 Glaucoma Symposium in Atlanta, Georgia, USA, by Y Liu.

